# A large proportion of poor birth outcomes among Aboriginal Western Australians are attributable to smoking, alcohol and substance misuse, and assault

**DOI:** 10.1101/553065

**Authors:** Alison J. Gibberd, Judy M. Simpson, Jocelyn Jones, Robyn Williams, Fiona Stanley, Sandra J. Eades

## Abstract

**Background:** Aboriginal infants have poorer birth outcomes than non-Aboriginal infants. Harmful use of tobacco, alcohol, and other substances is higher among Aboriginal women, as is violence, due to factors such as intergenerational trauma and poverty. We estimated the proportion of small for gestational age (SGA) births, preterm births, and perinatal deaths that could be attributed to these risks.

**Methods:** Birth, hospital, mental health, and death records for Aboriginal singleton infants born in Western Australia from 1998-2010 and their parents were linked. Using logistic regression with a generalized estimating equation approach, associations with birth outcomes and population attributable fractions were estimated after adjusting for demographic factors and maternal health during pregnancy.

**Results:** Of 28,119 births, 16% of infants were SGA, 13% were preterm, and 2% died perinatally. 51% of infants were exposed *in utero* to at least one of the risk factors and the fractions attributable to them were 37% (SGA), 16% (preterm) and 20% (perinatal death).

**Conclusions:** A large proportion of adverse outcomes were attributable to the modifiable risk factors of substance use and assault. Significant improvements in Aboriginal perinatal health are likely to follow reductions in these risk factors. These results highlight the importance of identifying and implementing risk reduction measures which are effective in, and supported by, Aboriginal women, families, and communities.

## Manuscript

### Background

Australian Aboriginal and Torres Strait Islander (hereafter respectfully referred to as Aboriginal) infants tend to have poorer birth outcomes than non-Aboriginal infants. In the state of Western Australia (WA), preterm birth, stillbirth, and neonatal death rates are 2-3 times higher and the average birthweight is 200g less for infants with Aboriginal mothers than non-Aboriginal mothers [1,2]. In the past three decades, though the neonatal death rate has declined, the rates of small for gestational age (SGA) births, preterm births and stillbirths to Aboriginal mothers in WA have remained static [1-3] and a better understanding of why these outcomes are so common among Aboriginal infants is needed.

Smoking during pregnancy, harmful use of alcohol and drugs, and assault against the mother are all associated with poor birth outcomes [4,5] and are also more common among Aboriginal than non-Aboriginal women. The context in which they arise is generations of displacement from traditional lands, limited education and employment opportunities resulting in economic disadvantage, marginalisation, racism, forced removal of children from their parents, and other associated losses.

Aboriginal women smoke during approximately half of all pregnancies [6]. While abstinence from alcohol is common in Aboriginal communities, among those who do drink, consumption is more likely to be harmful and Aboriginal women are seven times more likely to die from alcoholic liver cirrhosis and alcohol dependence than non-Aboriginal women [7]. Rates of drug use are also high. In 2015, 27% of Aboriginal women reported using substances in the previous 12 months for non-medical reasons [8]. By contrast, 13% of non-Aboriginal women reported illicit drug use [9]. Finally, Aboriginal women are 35 times more likely to be hospitalised because of an assault than non-Aboriginal women [10]. The use of tobacco, alcohol and drugs, and assault are inter-related and multiple risk factors can aggregate in pregnancy [11].

The associations of poor birth outcomes with smoking, alcohol, drugs, and assault have been observed across a range of populations [4,5]. However, their contribution to the high levels of poor birth outcomes among Aboriginal infants is rarely quantified, particularly for assault. We therefore aimed to estimate the proportions of Aboriginal SGA births, preterm births, and perinatal deaths in WA from 1998 to 2010 that can be attributed to smoking, misuse of alcohol or drugs, and assault.

### Methods

#### Study cohort and data sources

The study cohort comprised all singleton births in WA from 1998 to 2010, where the infant and their full siblings were categorised as Aboriginal using the algorithm MSM+Family, described in Gibberd *et al* [12]. Briefly, this algorithm assigns Aboriginal status to each infant using the Indigenous identifiers on their birth record (Midwives Notification System [MNS]), birth registration, inpatient hospital records (Hospital Morbidity Data Collection), and WA Register of Developmental Anomalies (WARDA) record, as well as their family members’ records. The algorithm offers some protection against false positives that can occur with linkage of many records, while relatives’ information resolves some false negatives and positives and reduces the number of infants with unknown Aboriginal status [12]. The study cohort’s relatives were identified by WA’s records of family links, the Family Connections System [13].

The Data Linkage Branch in the Department of Health linked the above datasets and death records using probabilistic linkage.

In total, 28,119 Aboriginal infants were recorded in the MNS from 1998 to 2010. Each birth with a gestational age of at least 20 weeks and/or a birthweight of 400 grams is notified to the WA Department of Health by an attending midwife or medical officer. Details of the mother and infant, the birth, and conditions affecting the mother or pregnancy are recorded on the birth record.

#### Outcomes

The three outcomes of interest were SGA, preterm birth, and perinatal death. An infant was defined as SGA if their birthweight was less than the first decile for Australian singleton infants of the same sex and gestational age, born alive from 1998 to 2007 [14]. Preterm birth was defined as any live birth or stillbirth at 20 to 36 completed weeks’ gestation. In line with the Australian policy of classification of perinatal deaths, they were defined as either stillbirth (the death of a baby prior to the complete expulsion or extraction from its mother at a gestational age of 20 or more completed weeks or with a birthweight of at least 400 grams) or death less than 28 days after a live birth [15].

Gestational age (GA) as determined by Blair *et al*’s method [16] was missing for 67 of the 28,119 infants. All 67 infants had an estimated gestational age of 20 to 34 weeks in their birth records, which was based on observations of the neonate, including sole creases and scalp hair. We classified all 67 infants as preterm because, even if this estimate was less accurate than Blair *et al*’s method, the magnitude of the error would need to be at least 3 weeks for an infant to be misclassified as preterm. However, as classification as SGA was based on actual GA, infants with missing GA were excluded from analyses involving SGA.

#### Study factors

The four risk factors of interest were maternal smoking, alcohol misuse, drug misuse, and assault during pregnancy. Maternal smoking has been recorded comprehensively on WA birth records since 1998. Mothers were categorised as misusing alcohol or drugs if an alcohol- or drug-related diagnosis was recorded in (1) the child’s birth record or (2) in any diagnosis field on their hospital admissions during the pregnancy or (3) a mental health record during the pregnancy.

The mother was categorised as the victim of assault if violence against her was recorded in a hospital admission in the period from two years prior to the start of the pregnancy until the birth. We included this ‘look-back’ period because around half of all women who suffer physical violence before pregnancy continue to be exposed during pregnancy and socio-economically deprived women are particularly likely to have violence continue into pregnancy [17], violence prior to pregnancy is an independent predictor of poor birth outcomes [18], and a look-back period was likely to improve ascertainment of cases of assault as we could only identify violence through external injury codes in one dataset. We did not include a look-back period for alcohol or drug misuse as we had additional data sources to identify misuse and the vast majority of women cease or reduce their consumption during pregnancy [19, 20]. These issues did not arise for smoking, a mandatory field in the birth record.

#### Other explanatory variables

We initially chose explanatory variables that are known, or suspected, to be associated with the birth outcomes, were recorded in our data, and were not rare. They were *demographic information* (infant year of birth, infant sex, maternal parity, and maternal age), *maternal infections during pregnancy* (urinary tract infections [UTI], *Herpes simplex*, gonorrhoea, chlamydia, vaginitis [vaginitis, *Candida*, and/or trichomoniasis], Group B streptococcus, and other infections [syphilis, toxoplasmosis, rubella, cytomegalovirus, and/or varicella zoster]), and *maternal long-term health* (maternal height, diabetes, hypertension, obesity, mental health conditions, heart disease, and asthma).

We identified maternal health conditions from both the infant’s birth record and the mother’s hospital records during pregnancy as the combination improves ascertainment with little change in the number of false positives [21]. Using broad categories for diseases during pregnancy also increases sensitivity, with minimal change in specificity [21]. “Diabetes” included pre-existing and gestational diabetes, and “hypertension” included pre-existing hypertension complicating pregnancy, pre-eclampsia, and eclampsia. All relevant codes of the International Classification of Diseases, 9^th^ and 10^th^ Revisions (ICD-9-CM, ICD-10-AM) are listed in Additional file 1.

#### Analysis

In the adjusted logistic regression model for each outcome, we initially included the study factors (maternal smoking, alcohol misuse, drug misuse, assault) and all other explanatory variables with P < 0.2 in unadjusted models. Variables were sequentially removed until only variables with P < 0.05 and the study factors remained. We then entered interactions with the study factors, except interactions with maternal height which we did not believe were biologically plausible. With no prior reason to believe there would be interactions, we set the significance level at 0.01. We then entered all significant interactions into the model simultaneously and retained those that remained significant. We then checked the variable selection by adding the excluded variables to the model again, one-by-one. However, they remained non-significant and their inclusion did not meaningfully change the coefficients for the four study factors.

We used the multivariable fractional polynomial procedure to test whether non-linear functional forms for the continuous variables were preferable [22]. In the fully adjusted models, maternal height had linear associations with all three outcomes and infant’s year of birth had linear or no association. Transformations of maternal age of degree 1 and degree 3 were selected for the preterm birth model and degree -½ and degree 3 for the perinatal death model.

Maternal height, an important predictor of birth outcomes [23], was missing for 9305/28119 (33%) of births. However, for 6078/9305 cases, maternal height was available in siblings’ birth records. We used multiple imputation to impute the remaining 3227 (11%) missing cases [Additional file 1] [24]. We created 20 complete datasets.

Regression coefficients and variances were obtained from models fit to each of the 20 datasets using logistic regression with a generalised estimating equation (GEE) approach to account for correlation within mothers. Independent working correlation matrices and robust standard errors were selected. Using Rubin’s rules, we combined the 20 sets of regression coefficients and variances [25].

Parents may have children with more than one partner and those partners may also have children with more than one partner. As a result, children are not all clustered in nuclear families and can be cross-classified to mothers and fathers. We calculated regression coefficient covariance matrices that took cross-classification into account [Additional file 1]. Because these matrices were very similar to those obtained by clustering on the mother, we present the results from clustering by mother only.

Population attributable fractions (PAFs) are the proportions of disease attributable to an exposure or group of exposures. We calculated model-based adjusted PAFs for the risk factors of interest by calculating the difference between the observed number of poor outcomes and the expected number if the risk factor was eliminated from the population, divided by the observed number of outcomes [26]. We estimated 95% confidence intervals using bootstrap with 1,000 replicates.

SAS software, Version 9.4, was used for all analyses, with some exceptions. R 3.4.0 [27] was used for the multiple imputation, to identify appropriate fractional polynomials, to obtain bootstrap samples, and to calculate population attributable fractions (using regression coefficients obtained using SAS).

#### Sensitivity and subgroup analyses

We conducted sensitivity analyses by analysing the subset of 18,814 out of 28,119 births which had maternal height recorded on their birth records (‘complete cases’). As we did not include remoteness or socioeconomic disadvantage to avoid overfitting, sensitivity analyses were also conducted with these variables included. Finally, as research often focuses on first-born infants and birth weight varies by parity, we also stratified by parity.

### Results

Approximately a quarter (27%) of the 28,119 infants had at least one of the three outcomes of interest: 16% of infants were SGA; 13% of infants were preterm; and 2% died perinatally (Table 1). Mothers smoked during 47% of the pregnancies and alcohol misuse was recorded for 3% of pregnancies, drug misuse for 6%, and assault for 7%. For 51% of births, at least one of these risks was present.

**Table 1:**
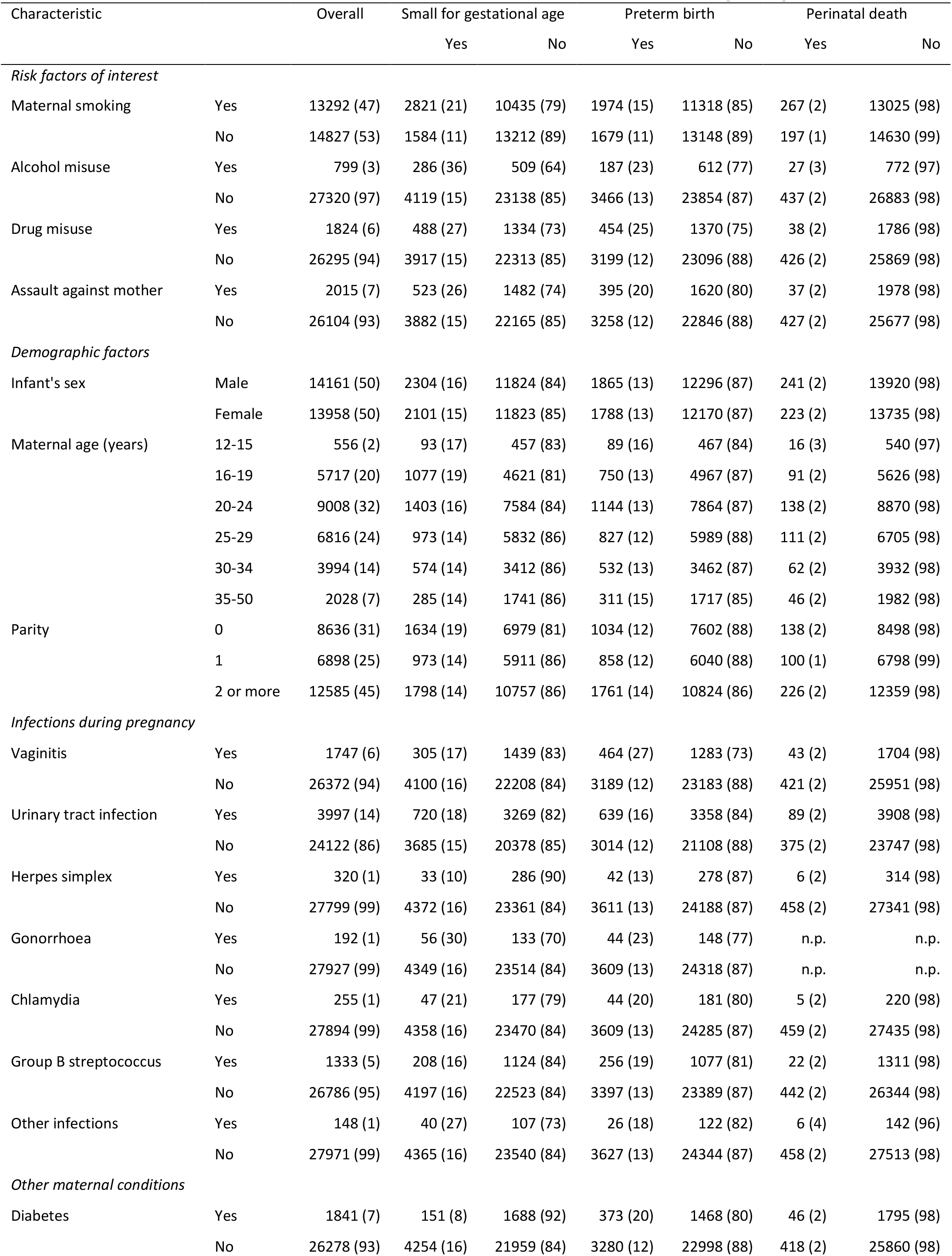
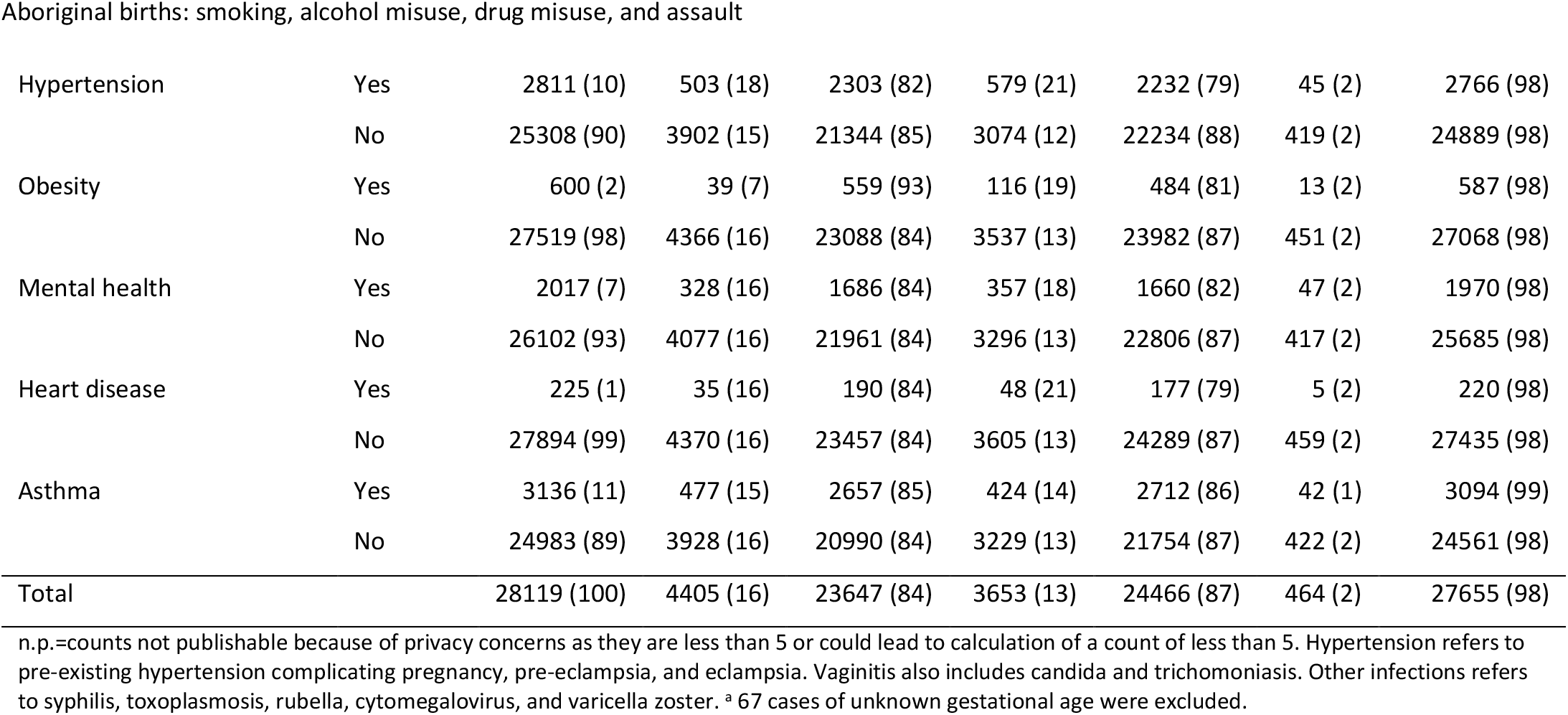
Associations between birth outcomes and infant and maternal characteristics for 28,119 WA Aboriginal singletons born 1998-2010

| Characteristic |  | Overall | Small for gestational age |  | Preterm birth |  | Perinatal death |  |
| --- | --- | --- | --- | --- | --- | --- | --- | --- |
|  |  |  | Yes | No | Yes | No | Yes | No |
| *Risk factors of interest* |  |  |  |  |  |  |  |  |
| Maternal smoking | Yes | 13292 (47) | 2821 (21) | 10435 (79) | 1974 (15) | 11318 (85) | 267 (2) | 13025 (98) |
|  | No | 14827 (53) | 1584 (11) | 13212 (89) | 1679 (11) | 13148 (89) | 197 (1) | 14630 (99) |
| Alcohol misuse | Yes | 799 (3) | 286 (36) | 509 (64) | 187 (23) | 612 (77) | 27 (3) | 772 (97) |
|  | No | 27320 (97) | 4119 (15) | 23138 (85) | 3466 (13) | 23854 (87) | 437 (2) | 26883 (98) |
| Drug misuse | Yes | 1824 (6) | 488 (27) | 1334 (73) | 454 (25) | 1370 (75) | 38 (2) | 1786 (98) |
|  | No | 26295 (94) | 3917 (15) | 22313 (85) | 3199 (12) | 23096 (88) | 426 (2) | 25869 (98) |
| Assault against mother | Yes | 2015 (7) | 523 (26) | 1482 (74) | 395 (20) | 1620 (80) | 37 (2) | 1978 (98) |
|  | No | 26104 (93) | 3882 (15) | 22165 (85) | 3258 (12) | 22846 (88) | 427 (2) | 25677 (98) |
| *Demographic factors* |  |  |  |  |  |  |  |  |
| Infant's sex | Male | 14161 (50) | 2304 (16) | 11824 (84) | 1865 (13) | 12296 (87) | 241 (2) | 13920 (98) |
|  | Female | 13958 (50) | 2101 (15) | 11823 (85) | 1788 (13) | 12170 (87) | 223 (2) | 13735 (98) |
| Maternal age (years) | 12-15 | 556 (2) | 93 (17) | 457 (83) | 89 (16) | 467 (84) | 16 (3) | 540 (97) |
|  | 16-19 | 5717 (20) | 1077 (19) | 4621 (81) | 750 (13) | 4967 (87) | 91 (2) | 5626 (98) |
|  | 20-24 | 9008 (32) | 1403 (16) | 7584 (84) | 1144 (13) | 7864 (87) | 138 (2) | 8870 (98) |
|  | 25-29 | 6816 (24) | 973 (14) | 5832 (86) | 827 (12) | 5989 (88) | 111 (2) | 6705 (98) |
|  | 30-34 | 3994 (14) | 574 (14) | 3412 (86) | 532 (13) | 3462 (87) | 62 (2) | 3932 (98) |
|  | 35-50 | 2028 (7) | 285 (14) | 1741 (86) | 311 (15) | 1717 (85) | 46 (2) | 1982 (98) |
| Parity | 0 | 8636 (31) | 1634 (19) | 6979 (81) | 1034 (12) | 7602 (88) | 138 (2) | 8498 (98) |
|  | 1 | 6898 (25) | 973 (14) | 5911 (86) | 858 (12) | 6040 (88) | 100 (1) | 6798 (99) |
|  | 2 or more | 12585 (45) | 1798 (14) | 10757 (86) | 1761 (14) | 10824 (86) | 226 (2) | 12359 (98) |
| *Infections during pregnancy* |  |  |  |  |  |  |  |  |
| Vaginitis | Yes | 1747 (6) | 305 (17) | 1439 (83) | 464 (27) | 1283 (73) | 43 (2) | 1704 (98) |
|  | No | 26372 (94) | 4100 (16) | 22208 (84) | 3189 (12) | 23183 (88) | 421 (2) | 25951 (98) |
| Urinary tract infection | Yes | 3997 (14) | 720 (18) | 3269 (82) | 639 (16) | 3358 (84) | 89 (2) | 3908 (98) |
|  | No | 24122 (86) | 3685 (15) | 20378 (85) | 3014 (12) | 21108 (88) | 375 (2) | 23747 (98) |
| Herpes simplex | Yes | 320 (1) | 33 (10) | 286 (90) | 42 (13) | 278 (87) | 6 (2) | 314 (98) |
|  | No | 27799 (99) | 4372 (16) | 23361 (84) | 3611 (13) | 24188 (87) | 458 (2) | 27341 (98) |
| Gonorrhoea | Yes | 192 (1) | 56 (30) | 133 (70) | 44 (23) | 148 (77) | n.p. | n.p. |
|  | No | 27927 (99) | 4349 (16) | 23514 (84) | 3609 (13) | 24318 (87) | n.p. | n.p. |
| Chlamydia | Yes | 255 (1) | 47 (21) | 177 (79) | 44 (20) | 181 (80) | 5 (2) | 220 (98) |
|  | No | 27894 (99) | 4358 (16) | 23470 (84) | 3609 (13) | 24285 (87) | 459 (2) | 27435 (98) |
| Group B streptococcus | Yes | 1333 (5) | 208 (16) | 1124 (84) | 256 (19) | 1077 (81) | 22 (2) | 1311 (98) |
|  | No | 26786 (95) | 4197 (16) | 22523 (84) | 3397 (13) | 23389 (87) | 442 (2) | 26344 (98) |
| Other infections | Yes | 148 (1) | 40 (27) | 107 (73) | 26 (18) | 122 (82) | 6 (4) | 142 (96) |
|  | No | 27971 (99) | 4365 (16) | 23540 (84) | 3627 (13) | 24344 (87) | 458 (2) | 27513 (98) |
| *Other maternal conditions* |  |  |  |  |  |  |  |  |
| Diabetes | Yes | 1841 (7) | 151 (8) | 1688 (92) | 373 (20) | 1468 (80) | 46 (2) | 1795 (98) |
|  | No | 26278 (93) | 4254 (16) | 21959 (84) | 3280 (12) | 22998 (88) | 418 (2) | 25860 (98) |
| Hypertension | Yes | 2811 (10) | 503 (18) | 2303 (82) | 579 (21) | 2232 (79) | 45 (2) | 2766 (98) |
|  | No | 25308 (90) | 3902 (15) | 21344 (85) | 3074 (12) | 22234 (88) | 419 (2) | 24889 (98) |
| Obesity | Yes | 600 (2) | 39 (7) | 559 (93) | 116 (19) | 484 (81) | 13 (2) | 587 (98) |
|  | No | 27519 (98) | 4366 (16) | 23088 (84) | 3537 (13) | 23982 (87) | 451 (2) | 27068 (98) |
| Mental health | Yes | 2017 (7) | 328 (16) | 1686 (84) | 357 (18) | 1660 (82) | 47 (2) | 1970 (98) |
|  | No | 26102 (93) | 4077 (16) | 21961 (84) | 3296 (13) | 22806 (87) | 417 (2) | 25685 (98) |
| Heart disease | Yes | 225 (1) | 35 (16) | 190 (84) | 48 (21) | 177 (79) | 5 (2) | 220 (98) |
|  | No | 27894 (99) | 4370 (16) | 23457 (84) | 3605 (13) | 24289 (87) | 459 (2) | 27435 (98) |
| Asthma | Yes | 3136 (11) | 477 (15) | 2657 (85) | 424 (14) | 2712 (86) | 42 (1) | 3094 (99) |
|  | No | 24983 (89) | 3928 (16) | 20990 (84) | 3229 (13) | 21754 (87) | 422 (2) | 24561 (98) |
| Total |  | 28119 (100) | 4405 (16) | 23647 (84) | 3653 (13) | 24466 (87) | 464 (2) | 27655 (98) |

n.p.=counts not publishable because of privacy concerns as they are less than 5 or could lead to calculation of a count of less than 5. Hypertension refers to pre-existing hypertension complicating pregnancy, pre-eclampsia, and eclampsia. Vaginitis also includes candida and trichomoniasis. Other infections refers to syphilis, toxoplasmosis, rubella, cytomegalovirus, and varicella zoster. <sup>a</sup> 67 cases of unknown gestational age were excluded.

Maternal smoking was associated with over twice the odds of SGA birth, 26% higher odds of preterm birth and 49% higher odds of perinatal death (Figure 1). Alcohol was associated with 118% higher odds of SGA and 83% high odds of perinatal death, but the association with preterm birth, while positive, was not statistically significant. Drug misuse and assault were strongly associated with SGA and preterm birth, but not perinatal death.

**Figure 1:**
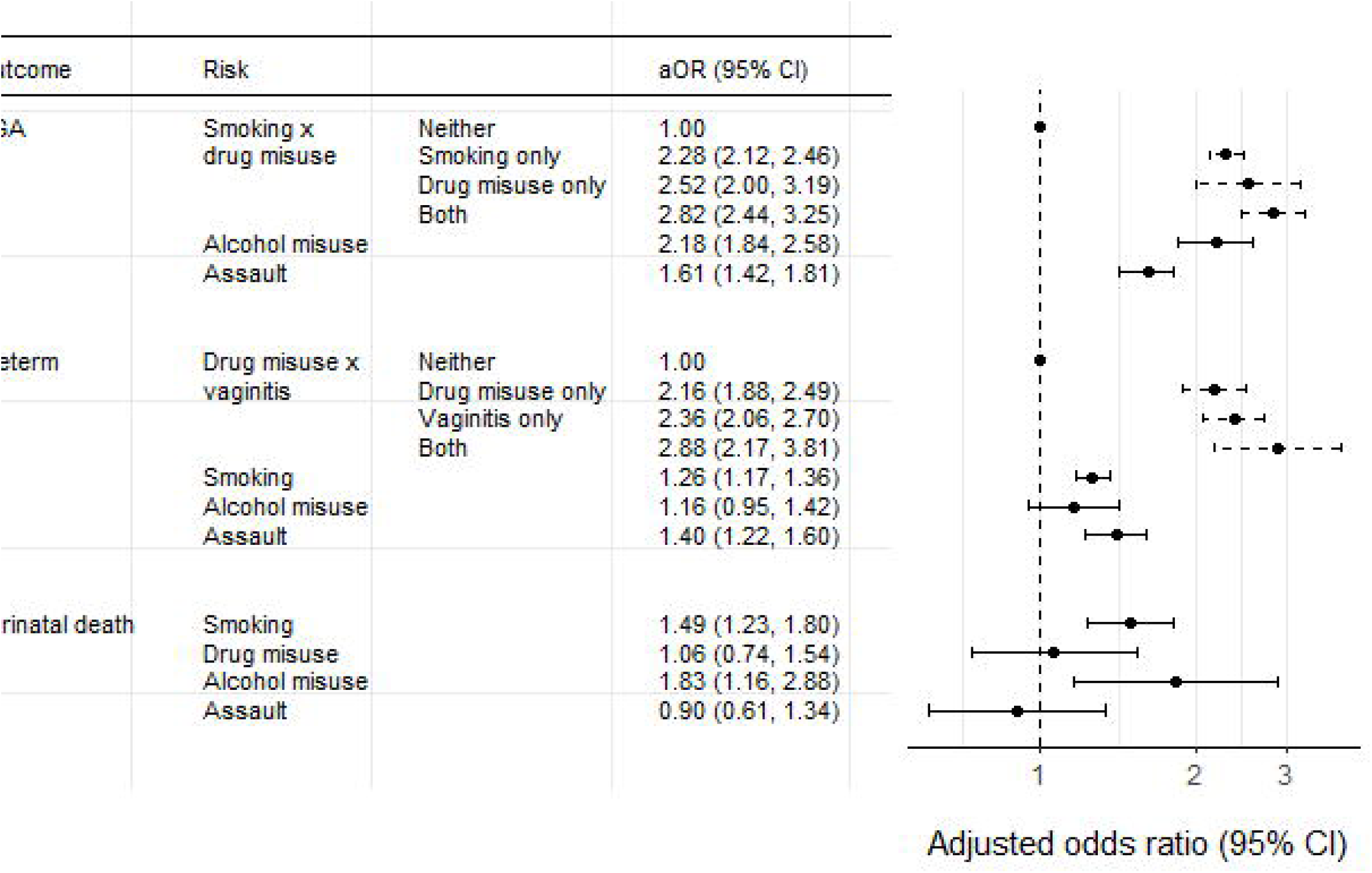
Adjusted odds ratios (aOR) of birth outcomes from smoking, alcohol misuse, drug misuse, and assault. Adjusted odds ratios are for 28,119 Aboriginal singleton infants born in Western Australia, 1998-2010. Bars are 95% confidence intervals. Each model adjusted for maternal smoking, drug misuse, alcohol misuse, assault, maternal height and diabetes. The model for SGA also included an interaction between maternal smoking and drug misuse, infant sex, parity, hypertension (pre-existing hypertension complicating pregnancy, pre-eclampsia, and eclampsia), obesity, gonorrhoea, herpes, and other infections (syphilis, toxoplasmosis, rubella, cytomegalovirus, and varicella zoster). The model for preterm birth also included an interaction between drug misuse and vaginitis (vaginitis, candida and trichomoniasis), maternal age, parity, infant’s year of birth, hypertension, heart disease, urinary tract infection, Group B streptococcus, obesity, mental health conditions, and gonorrhoea. The model for perinatal death also included maternal age and urinary tract infection. Confidence intervals are dashed for risks with interactions and solid otherwise.

There were two interactions with the risk factors of interest. Compared to mothers who neither smoked nor misused drugs, those who either smoked or misused drugs had over twice the odds of a SGA infant (adjusted odds ratio (aOR) 2.28 [95% CI: 2.12, 2.46] for smoking only and 2.52 [95% CI: 2.00, 3.19] for drug misuse only). However, if the mother both smoked and misused drugs, the infant’s odds of being SGA were not much greater (aOR 2.82 [95% CI: 2.44, 3.25]). Similarly, for preterm birth, there was an interaction between drug misuse and vaginitis.

37% of SGA births, 16% of preterm births and 20% of perinatal deaths could be attributed to smoking, alcohol misuse, drug misuse or assault. As PAFs are affected by prevalence of the risk factor, as well as the magnitude of the risk, smoking had the highest PAF for each outcome (Figure 2).

**Figure 2:**
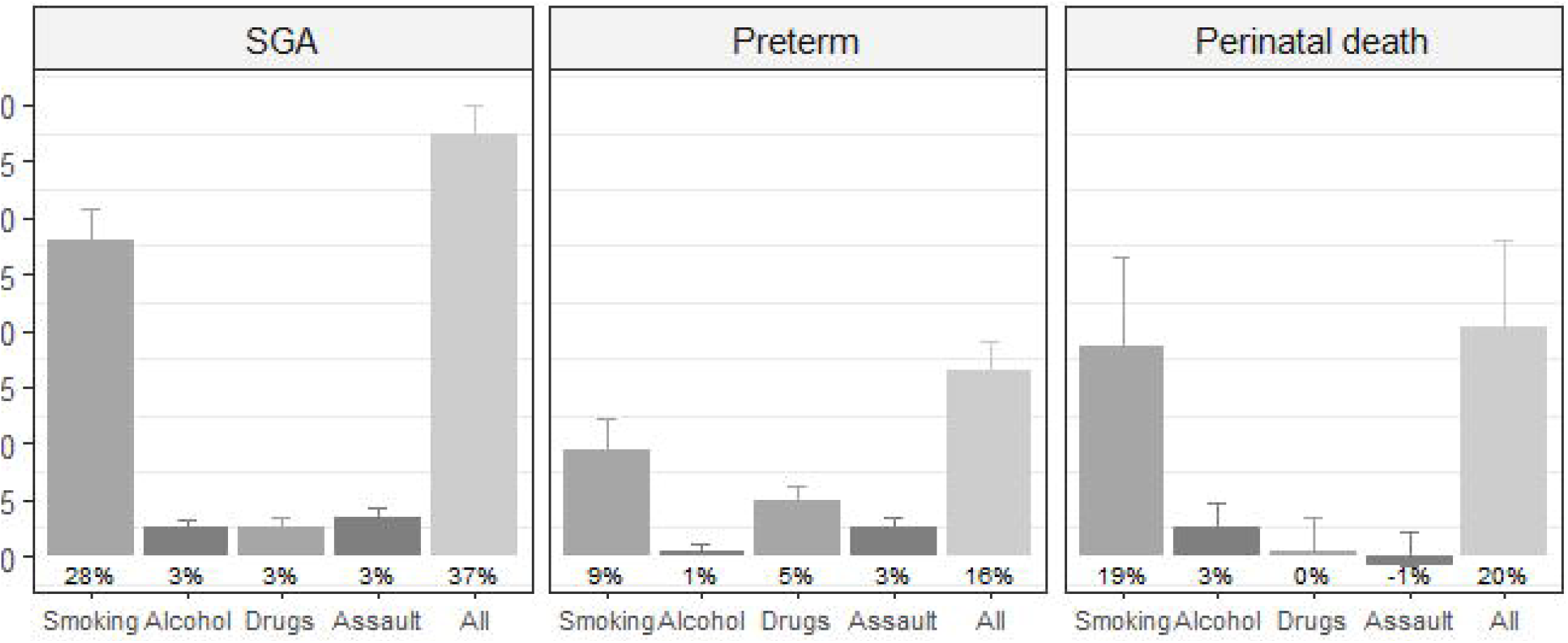
Adjusted population attributable fractions for birth outcomes from smoking, alcohol misuse, drug misuse, and assault. Adjusted population attributable fractions with 95% confidence intervals are for 28,119 Aboriginal singleton infants born in Western Australia, 1998-2010. Each model adjusted for maternal smoking, drug misuse, alcohol misuse, assault, maternal height and diabetes. The model for SGA also included an interaction between maternal smoking and drug misuse, infant sex, parity, hypertension (pre-existing hypertension complicating pregnancy, pre-eclampsia, and eclampsia), obesity, gonorrhoea, herpes, and other infections (syphilis, toxoplasmosis, rubella, cytomegalovirus, and varicella zoster). The model for preterm birth also included an interaction between drug misuse and vaginitis (vaginitis, candida and trichomoniasis), maternal age, parity, infant’s year of birth, hypertension, heart disease, urinary tract infection, Group B streptococcus, obesity, mental health conditions, and gonorrhoea. The model for perinatal death also included maternal age and urinary tract infection.

Results from analyses of complete cases were similar to the main results using imputed maternal heights, with the exception of perinatal death, where the odds were only 29% higher for alcohol misuse than no misuse among the complete cases, compared to 83% higher among the full sample [Tables 6-8 in Additional file 1]. The addition of remoteness and socioeconomic disadvantage to the models slightly attenuated the significant relationships between the poor birth outcomes and alcohol misuse and assault. However, the odds ratios for smoking changed little and those for drug misuse increased. The PAFs changed little with the inclusion of remoteness and socioeconomic disadvantage [Table 9 in Additional file 1].

### Discussion

Between 1998 and 2010, 27% of all Aboriginal infants in WA were SGA, preterm, or died perinatally. A substantial proportion of these outcomes could be attributed to *in utero* exposure to maternal smoking, alcohol misuse, drug misuse, and assault against their mother – 37% of SGA births, 16% of preterm births, and 20% of perinatal deaths. With half (51%) of the infants exposed to at least one of these risk factors, reductions in these interrelated behaviours may greatly improve Aboriginal perinatal health.

More poor outcomes could be attributed to smoking than the other risks, reflecting the fact that 47% of infants were exposed. We found 28% of SGA births could be attributed to smoking. By contrast, Taylor *et al* found that for infants born to mothers in the state of New South Wales (NSW, 97% non-Aboriginal mothers) with a smoking rate of 11% during pregnancy, only 10% of SGA births for term infants with non-diabetic mothers were attributable to smoking, 3% for term infants with diabetic mothers, and 12% of SGA births for preterm infants [28].

It is likely we underestimated the prevalence of alcohol and drug misuse and assault as this information was not mandatory in the datasets. It is also likely that we identified the more serious cases, given we identified most cases through hospital admissions and the nature of ICD diagnoses. Our estimates may also be lower than other studies as we included Aboriginal infants with non-Aboriginal mothers (18% of births). For example, only 0.5% of infants to non-Aboriginal mothers were categorised as exposed to alcohol, compared to 3.3% of infants with Aboriginal mothers.

The true proportion of infants subject to harmful levels of alcohol *in utero* is difficult to assess. It is not clear which drinking patterns (timing, frequency, and amount) are harmful and few studies have collected detailed information from Aboriginal women. In WA studies, 23% or more Aboriginal women drank during pregnancy, though this included any alcohol consumption [29], harmful consumption from age 10 to a year after pregnancy [30], or the sample was highly selected, including a community with high average consumption [31, 32].

Our finding that 6% of infants were exposed to drug misuse may be more accurate than our finding about alcohol misuse. WA Aboriginal mothers have reported using marijuana during pregnancy for 9% of births and other drugs for fewer than 1% [29], though some studies from other states have found greater drug use [11, 33].

The proportion of infants whose mothers were categorised as victims of assault (7%) was half the proportion of all Aboriginal women reported by the Australian Bureau of Statistics to have experienced physical violence in the past year in 2014-15 (14%) [34]. Nevertheless, the higher odds of SGA and preterm birth following assault, compared to no assault, (aOR: 1.61 and 1.40, respectively) were similar to those from a 2010 meta-analysis for low birth weight (aOR: 1.53 [95% CI: 1.28, 1.82]) and preterm birth (1.46 [95% CI: 1.27, 1.67]) [4]. Hypotheses about the effect of maternal stress hormones during pregnancy have been proposed, for example, the release of oxytocin could induce early contractions [35].

Although we underestimated the prevalence of some risk factors, as they are often clustered in women the PAF for all risks combined may be a reasonable estimate of the true PAF. For example, if a woman smoked and misused alcohol, but only smoking was documented, her contribution to the risk estimate for smoking may encompass the effects of both smoking and alcohol, resulting in a higher risk estimate for smoking. When the PAF was calculated for all risk factors combined, some of this additional risk from (unidentified) alcohol misuse would be captured in the combined PAF.

Approximately 1% of pregnant Aboriginal women in WA have no antenatal care and 17% have no care until after 24 weeks’ gestation [2]. Antenatal care attendance could not be explored in our analysis as this information was not available for the birth years covered by this study. Substance misuse and assault and suboptimal antenatal care are clearly associated. However, it is not clear whether substance misuse and assault affect attendance for antenatal care or whether they share common causes. If the former, antenatal care is an intermediate variable and including antenatal care in the model might have biased the results. If the latter, and if the common causes are not adjusted for, our estimates of the effects of the risk factors may be biased upwards.

Births following terminations of pregnancy from 20 weeks’ gestation or with congenital abnormalities were not excluded because we were interested in population-level outcomes, substance misuse is associated with certain developmental abnormalities [36], and some of the poor birth outcomes in this study may have followed the combination of developmental abnormalities and the risk factors of interest. The proportion of preventable poor birth outcomes would have been higher in a sample of births which did not include terminations of pregnancy or congenital abnormalities than in the full population, most likely resulting in higher estimates of the PAFs.

For this study we used a composite endpoint of stillbirths and neonatal deaths, as estimates obtained from modelling the 141 neonatal deaths separately would have a high degree of uncertainty, ‘live-birth bias’ may bias our estimates, and because the causes of the majority of neonatal deaths arise prenatally or in the intrapartum period [37, 38].

Smoking, alcohol and drug misuse, and violence are intrinsically linked and may be triggered by boredom, unemployment, marginalisation, poor mental health, overcrowded housing, and other stresses more commonly experienced by Aboriginal people [39-41]. Many Aboriginal people have complex health and social needs and some mainstream initiatives have been less effective in Aboriginal populations. For example, while smoking among pregnant Aboriginal women has dropped considerably, the decrease in recent years has been greater for non-Aboriginal women and non-Aboriginal women are more likely to quit smoking during pregnancy [6].

Aboriginal women can face significant barriers to change. Smoking and violence are normalised in some communities [39, 42] and drinking is frequently social with 27% of 180 Nyoongar women reporting drinking with a male partner while pregnant (Robyn Williams, personal communication, 7 November 2017). Fear of losing children to child protection agencies can discourage women from seeking help with substance misuse and violence [40]. WA Aboriginal children are 17.5 times more likely to be in out-of-home care than non-Aboriginal children [43]. Numerous additional challenges may affect Aboriginal women, such as limited services [40, 41].

Despite the widespread acknowledgement that Aboriginal-specific risk reduction measures are needed, rigorous evaluations of Aboriginal-specific responses are rare [44-46]. To the best of our knowledge only one randomised clinical trial involving pregnant Aboriginal women has been conducted. This trial aimed to assess the effect of a smoking cessation intervention which included advice about quitting smoking at a woman’s first antenatal appointment, follow-up appointments with Aboriginal healthcare workers and midwives, and nicotine replacement therapy [47]. More women in the treatment (psychosocial) arm quit smoking than in the standard care arm, but the difference was not statistically significant. The trial faced difficulties with high staff turnover, possible contamination between the two arms, and over 30% loss to follow-up with only 176 completing. Methodologically rigorous studies can be more difficult in Indigenous populations, with challenges such as small sample sizes and funding time-limits may be insufficient to establish and maintain relationships with communities [48]. Funding incentives and alternative governance approaches may encourage more studies [48].

Evaluations of restrictions (including bans) on the supply of alcohol to communities have found that, with Aboriginal leadership and community support, they can be effective in reducing consumption and related harms like violence, despite some unintended consequences [49]. The 2018 decision to introduce a minimum price for a unit of alcohol across the NT may provide evidence of the impact of price signals [50].

The lack of evidence about how to prevent violence against Aboriginal women is unsurprising as, globally, little is known about what works [51]. Some Aboriginal communities have night patrols. Evidence of efficacy is limited but many patrols are valued by community members, local police, and service providers, suggesting they have a positive impact [40].

Relevant and accessible data are essential to improve the evidence base and calls have recently been made for increased data collection and linkage; for example, the establishment of a national data collection on violence and the expansion of perinatal data collections to include details of domestic violence and substance use [52, 53]. In WA, from 2017, the quantity and frequency of alcohol consumption during pregnancy will be available. Routinely-collected data can be a cost-effective way of evaluating programs. Outcomes can be passively measured at many time points, loss to follow up due to relocation within the state is minimized, and data collection may be more objective. With population-based data, robust estimates of the scale of these issues can be obtained, as in this study.

While the evidence-base for Aboriginal-specific risk reduction measures is limited, studies in other populations have identified effective approaches that could be tailored to Aboriginal communities.

An empirical evidence base is only one possible influence on health policy [46]. Evidence from other populations, studies of the acceptability and feasibility of interventions in an Aboriginal context, and the knowledge of community members and other stakeholders can inform risk reduction policies. There is widespread agreement that programs must be genuine partnerships or Aboriginal-led, tailored to local communities, holistic, targeted at the family and community level, as well as individual, and adequately supported [42, 54, 55]. A supportive, rather than punitive, approach towards Aboriginal women struggling with substance misuse and violence is needed.

Underlying smoking, alcohol and drug use, and violence in Aboriginal communities is a post-colonial history of dispossession, intergenerational trauma, structural racism, and poverty [41, 56]. Addressing the social determinants of these risk factors and poor mental health is an essential part of reducing these risks.

### Conclusions

With half of WA’s Aboriginal infants exposed *in utero* to the preventable risk factors of smoking, alcohol or drug misuse, or assault against the mother and a large proportion of poor birth outcomes attributable to this exposure, great improvements in the health of Aboriginal babies are possible with reductions in these risk factors. These results highlight the importance of identifying and implementing risk reduction measures which are effective in, and supported by, Aboriginal women, families and communities.

## Declarations

### Ethics approval and consent to participate

This study was approved by the Western Australian Aboriginal Health Ethics Committee (Ref 306 - 08/10) and the Western Australian Department of Health Ethics Committee (Ref 2010/42). Consent to participate was not required for this study.

### Consent for publication

Not applicable.

### Availability of data and material

The authors do not have permission from the data custodians to make available the data analysed in this study.

### Authors’ contributions

AG, JS, FS, and SE developed the research question. AG undertook the data analysis and wrote the first draft. JS, JJ, RW, and SE all supported the data analysis and commented on the drafts. SE obtained the funding and the data.

## Acknowledgements

We thank staff at the Western Australian Data Linkage Branch and the data custodians of the Midwives Notification System, Death Registrations, Hospital Morbidity Data Collection and Mental Health Information System for access to, and linkage of, the data. We would also like to thank staff at the Telethon Kids Institute for their involvement in the design of WAAIFS.

Additional file 1

## References

1. Farrant BM, Shepherd CC. Maternal ethnicity, stillbirth and neonatal death risk in Western Australia 1998-2010. Aust N Z J Obstet Gynaecol. 2016;56:532–536.

2. Hutchinson M, Joyce A. Western Australia’s Mothers and Babies, 2013: 31st Annual Report of the Western Australian Midwives’ Notification System: Perth (AUST): Department of Health, Western Australia; 2016.

3. Diouf I, Gubhaju L, Chamberlain C, McNamara B, Joshy G, Oats J, et al. Trends in maternal and newborn health characteristics and obstetric interventions among Aboriginal and Torres Strait Islander mothers in Western Australia from 1986 to 2009. Aust N Z J Obstet Gynaecol. 2016;56:245–251.

4. Shah PS, Shah J. Maternal exposure to domestic violence and pregnancy and birth outcomes: a systematic review and meta-analyses. J Womens Health. 2010;19:2017–2031.

5. Flenady V, Koopmans L, Middleton P, Froen JF, Smith GC, Gibbons K, et al. Major risk factors for stillbirth in high-income countries: a systematic review and meta-analysis. Lancet. 2011;377:1331–1340.

6. Australian Institute of Health and Welfare. Birthweight of babies born to Indigenous mothers. Canberra (AUST): AIHW; 2014.

7. Chikritzhs T, Liang W. Does the 2008 NATSISS underestimate the prevalence of high risk Indigenous drinking? In: Hunter B, Biddle N, editors. Survey Analysis for Indigenous Policy in Australia: Social Science Perspectives. Canberra (AUST): ANU E Press; 2012; p. 49–64.

8. Australian Bureau of Statistics. National Aboriginal and Torres Strait Islander Social Survey, 2014-15. Cat. No. 4714.0. 2016. http://www.abs.gov.au/ausstats/abs@.nsf/Lookup/by%20Subject/4714.0∼2014-15∼Main%20Features∼Health%20risk%20factors∼12. Accessed 28 Jul 2017.

9. Australian Institute of Health and Welfare. National Drug Strategy Household Survey 2016: detailed findings. Drug Statistics series no. 31. Cat. no. PHE 214. Canberra (AUST): AIHW; 2017.

10. Australian Health Ministers’ Advisory Council. Aboriginal and Torres Strait Islander Health Performance Framework Report 2010. Canberra (AUST): AHMAC; 2011.

11. Passey ME, Sanson-Fisher RW, D’Este CA, Stirling JM. Tobacco, alcohol and cannabis use during pregnancy: clustering of risks. Drug Alcohol Depend. 2014;134:44–50.

12. Gibberd AJ, Simpson JM, Eades SJ. Use of family relationships improved consistency of identification of Aboriginal people in linked administrative data. J Clin Epidemiol. 2017;90:144–155.

13. Glasson EJ, de Klerk NH, Bass AJ, Rosman DL, Palmer LJ, Holman CD. Cohort profile: The Western Australian Family Connections Genealogical Project. Int J Epidemiol 2008;37:30e5.

14. Dobbins TA, Sullivan EA, Roberts CL, Simpson JM. Australian national birthweight percentiles by sex and gestational age, 1998-2007. Med J Aust. 2012;197:291–294.

15. Australian Institute of Health and Welfare. Perinatal deaths in Australia: 2013–2014. Canberra (AUST): AIHW; 2018

16. Blair E, Liu Y, Cosgrove P. Choosing the best estimate of gestational age from routinely collected population-based perinatal data. Paediatr Perinat Epidemiol. 2004;18(4):270–6.

17. Taillieu TL, Brownridge DA. Violence against pregnant women: Prevalence, patterns, risk factors, theories, and directions for future research. Aggress Violent Behav. 2010;15:14–35.

18. Silverman JG, Decker MR, Reed E, Raj A. Intimate partner violence victimization prior to and during pregnancy among women residing in 26 US states: associxations with maternal and neonatal health. Am J Obstet Gynecol. 2006;195:140–148.

19. Harrison PA, Sidebottom AC. Alcohol and drug use before and during pregnancy: an examination of use patterns and predictors of cessation. Matern Child Health J. 2009;13:386.

20. Powers JR, McDermott LJ, Loxton DJ, Chojenta CL. A prospective study of prevalence and predictors of concurrent alcohol and tobacco use during pregnancy. Matern Child Health J. 2013;17:76–84.

21. Lain SJ, Hadfield RM, Raynes-Greenow CH, Ford JB, Mealing NM, Algert CS, et al. Quality of data in perinatal population health databases: a systematic review. Med Care. 2012; 50:e7–20.

22. Royston P, Sauerbrei W. Multivariable model-building: a pragmatic approach to regression anaylsis based on fractional polynomials for modelling continuous variables. West Sussex (UK): John Wiley & Sons; 2008.

23. Han Z, Lutsiv O, Mulla S, McDonald SD. Maternal height and the risk of preterm birth and low birth weight: a systematic review and meta-analyses. J Obstet Gynaecol Can. 2012; 34:721–746.

24. van Buuren S. Flexible imputation of missing data. Boca Raton (FL): CRC press; 2012.

25. Rubin DB. Multiple imputation for nonresponse in surveys. New York (NY): Wiley; 1987.

26. Greenland S, Drescher K. Maximum likelihood estimation of the attributable fraction from logistic models. Biometrics. 1993; 49:865–872.

27. R Core Team. R: A language and environment for statistical computing. R Foundation for Statistical Computing, Vienna, Austria. 2015 and 2017. Available from: https://www.R-project.org/.

28. Taylor LK, Lee YYC, Lim K, Simpson JM, Roberts CL, Morris J. Potential prevention of small for gestational age in Australia: a population-based linkage study. BMC Pregnancy Childbirth. 2013;13:210.

29. Zubrick SR, Lawrence DM, Silburn SR, Blair E, Milroy H, Wilkes T, et al. The Western Australian Aboriginal Child Health Survey: The Health of Aboriginal Children and Young People. Perth (AUST): Telethon Institute for Child Health Research; 2004.

30. O’Leary CM, Halliday J, Bartu A, D’Antoine H, Bower C. Alcohol-use disorders during and within one year of pregnancy: a population-based cohort study 1985-2006. BJOG. 2013;120:744–753.

31. Eades SJ. Bibbulung Gnarneep (Solid Kid): A Longitudinal Study of a Population Based Cohort of Urban Aboriginal Children in Western Australia: Determinants of Health Outcomes During Early Childhood of Aboriginal Children Residing in an Urban Area. Perth (AUST): University of Western Australia; 2004.

32. Fitzpatrick JP, Latimer J, Ferreira ML, Carter M, Oscar J, Martiniuk AL, et al. Prevalence and patterns of alcohol use in pregnancy in remote Western Australian communities: The Lililwan Project. Drug Alcohol Rev. 2015;34:329–339.

33. Brown SJ, Mensah FK, Ah Kit J, Stuart-Butler D, Glover K, Leane C, et al. Use of cannabis during pregnancy and birth outcomes in an Aboriginal birth cohort: a cross-sectional, population-based study. BMJ Open. 2016;6:e010286.

34. Australian Bureau of Statistics. National Aboriginal and Torres Strait Islander Social Survey, 2014-15. Cat. No. 4714.0. 2016. http://www.abs.gov.au/AUSSTATS/abs@.nsf/DetailsPage/4714.02014-15?OpenDocument. Accessed 14 March 2018.

35. Graignic-Philippe R, Dayan J, Chokron S, Jacquet AY, Tordjman S. Effects of prenatal stress on fetal and child development: a critical literature review. Neurosci Biobehav Rev. 2014;43:137–162.

36. Viteri OA, Soto EE, Bahado-Singh RO, Christensen CW, Chauhan SP, Sibai BM. Fetal anomalies and long-term effects associated with substance abuse in pregnancy: a literature review. Am J Perinatol. 2015;32(5):405–16.

37. Liew Z, Olsen J, Cui X, Ritz B, Arah OA. Bias from conditioning on live birth in pregnancy cohorts: an illustration based on neurodevelopment in children after prenatal exposure to organic pollutants. Int J Epidemiol. 2015;44(1):345–54.

38. Australian Institute of Health and Welfare. Perinatal deaths in Australia: 2013–2014. Canberra (AUST): AIHW; 2018

39. Wood L, France K, Hunt K, Eades S, Slack-Smith L. Indigenous women and smoking during pregnancy: knowledge, cultural contexts and barriers to cessation. Soc Sci Med. 2008;66:2378–2389.

40. Blagg H, Bluett-Boyd N, Williams E. Innovative models in addressing violence against Indigenous women: State of knowledge paper. Sydney (AUST): ANROWS; 2015.

41. Gray D, Cartwright K, Stearne A, Saggers S, Wilkes E, Wilson M. Review of the harmful use of alcohol among Aboriginal and Torres Strait Islander people. Australian Indigenous HealthInfoNet. 2017.

42. Blagg H, Williams E, Cummings E, Hovane V, Torres M, Woodley KN. Innovative models in addressing violence against Indigenous women: Final report. Sydney (AUST): ANROWS; 2018.

43. SNAICC - National Voice for our Children. The Family Matters Report 2017: Measuring trends to turn the tide on the over-representation of Aboriginal and Torres Strait Islander children in out-of-home care in Australia. 2017.

44. Leske S, Harris MG, Charlson FJ, Ferrari AJ, Baxter AJ, Logan JM, et al. Systematic review of interventions for Indigenous adults with mental and substance use disorders in Australia, Canada, New Zealand and the United States. Aust N Z J Psychiatry. 2016;50:1040–1054.

45. Whitty M, Clifford A. The Dissemination of Alcohol Interventions for Indigenous Australians: A Mixed Studies Review Using Narrative Synthesis. J Alcohol Drug Depend. 2017;5:2.

46. Vujcich D, Rayner M, Allender S, Fitzpatrick R. When there is not enough evidence and when evidence is not enough: an Australian Indigenous smoking policy study. Front Public Health. 2016;4:228.

47. Eades SJ, Sanson-Fisher RW, Wenitong M, Panaretto K, D’Este C, Gilligan C, et al. An intensive smoking intervention for pregnant Aboriginal and Torres Strait Islander women: a randomised controlled trial. Med J Aust. 2012;197:42–46.

48. Paul CL, Sanson-Fisher R, Stewart J, Anderson AE. Being sorry is not enough: the sorry state of the evidence base for improving the health of indigenous populations. Am J Prev Med. 2010;38:566–568.

49. Muhunthan J, Angell B, Hackett ML, Wilson A, Latimer J, Eades A-M, et al. Global systematic review of Indigenous community-led legal interventions to control alcohol. BMJ Open. 2017; 7.

50. Northern Territory Government. Northern Territory Alcohol Harm Minimisation Plan 2018- 2019. Darwin (AUST): Northern Territory Government; 2018

51. Jahanfar S, Howard LM, Medley N. Interventions for preventing or reducing domestic violence against pregnant women. Cochrane Libr. 2014.

52. National Congress. Aboriginal and Torres Strait Islander Peak Organisations Unite: The Redfern Statement. 2016. https://nationalcongress.com.au/wp-content/uploads/2017/02/The-Redfern-Statement-9-June-_Final.pdf. Accessed 6 Mar 2018.

53. Australian Institute of Health and Welfare. Enhancing maternity data collection and reporting in Australia: National Maternity Data Development Project Stage 2. Cat. no. PER 73. Canberra (AUST): AIHW; 2016.

54. Upton P, Davey R, Evans M, Mikhailovich K, Simpson L, Hacklin D. Tackling Indigenous Smoking and Healthy Lifestyle Programme Review: A Multi-criteria Decision Analysis. Canberra (AUST): University of Canberra; 2014.

55. Intergovernmental Committee on Drugs. National Aboriginal and Torres Strait Islander peoples’ drug strategy 2014-2019: A sub-strategy of the National Drug Strategy 2010 - 2015. Canberra (AUST): Australian Government; 2015.

56. Ranzijn R, Nolan W. Psychology and Indigenous Australians: Foundations of cultural competence. Melbourne (AUST): Palgrave Macmillan Australia; 2009.

