## Supplementary material for "A large proportion of poor birth outcomes among Aboriginal Western Australians are attributable to smoking, alcohol and substance misuse, and assault"

**1. International Classification of Diseases codes**

To identify alcohol misuse, drug misuse, assault, and maternal health conditions, we used diagnoses on maternal hospital admissions, mental health records and her offspring’s birth record (Supplementary Table 1). Diagnoses are recorded using codes of the International Classification of Diseases, 9^th^ and 10^th^ Revisions (ICD-9-CM, ICD-10-AM) (ICD codes).

For the birth records, an attending midwife or medical officer records details of maternal health on a Notification of Case Attended (NOCA) form. Certain common conditions affecting the mother or pregnancy are recorded on the NOCA using check boxes. For other conditions, or free text (which is later transcribed to ICD codes on the birth record) are used [1].

**Supplementary Table 1: Codes of the International Classification of Diseases, 9^th^ and 10^th^ Revisions (ICD-9-CM, ICD-10-AM) used to identify alcohol and drug misuse, assault, infections during pregnancy, and other maternal health conditions**

|  | ICD-9-CM | ICD-10-AM |
| --- | --- | --- |
| **Alcohol misuse**^1^ | 255.0, 291, 303, 305.0, 331.7, 357.5, 359.4, 425.5, 535.30, 535.31, 571.0-571.3, 577.1, 655.43, 790.3, 980.0, 980.8, 980.9, E860.0, E860.9, E950.9, E980.5, V57.89, V65.42, V11.3 | E24.4, F10, G31.2, G62.1, G72.1, I42.6, K29.2, K70, K86.0, O35.4, R78.0, T51.0, T51.8, T51.9, X45, X65, Y15, Y90, Y91, Z72.1 |
| **Drug misuse**^2^ |  |  |
|  | 292, 304, 305.2-305.9, 648.3, 655.5, 965.0, 965.8, 967, 968.2, 968.3, 968.5, 969.3-969.9, 970.1, 970.8, 971.1, 977.0, E850.0-E850.2, E850.8, E851, E852, E853.2, E853.8, E853.9, E854.1-E854.3, E855.1, E855.2, E855.4, E858.8, E935.0, E980.0, E980.1, E980.3, E980.4, V65.42 | F11-F19, F55.0, F55.9, O35.5, R78.1-R78.5, T39.8, T40, T41.0, T41.1, T42.3, T42.4, T43.6, T43.9, T50.7, T50.9, X40-X42, X44, Y10, Y11, Y14, Z71.5, Z72.2 |
| **Assault** | E96 | X85-X99, Y00-Y09 |
| **Vaginitis** | 112.1, 112.2, 112.9, 131, 616.1 | A59, B37.3, B37.4, B37.9, N76.0, N76.1, N76.2, N76.3, N77.1 |
| **Urinary tract infection** | 590, 595, 599.0 | N10-N12, N30, N39.0, O23.0-O23.4 |
| ***Herpes simplex*** | 054 | A60, B00 |
| **Chlamydia** | 077.98, 078.88, 079.88, 099.1, 099.41, 099.5 | A55, A56, A74 |
| **Gonorrhoea** | 098, 647.1, V02.7 | A54, O98.2 |
| **Group B streptococcus** | 041.0, 041.1 | B95 |
| **Other infections**^3^ | 052, 053, 056, 078.5, 091-097, 130, 484.1, 647.0, 647.5 | A51-A53, B01, B02, B06, B25, B27.1, B58, O98.1 |
| **Diabetes** | 250, 648.0, 648.8 | E10-E14, O24 |
| **Hypertension** | 642 | O10, O11, O13-O16 |
| **Obesity** | 278.0 | E66 |
| **Mental health**^4^ | 290, 293-302, 306-319, E950-E959 | F00-F09, F20-F99, X60-X84 |
| **Heart disease** | 393-398, 428 | I05-I09, I11.0, I13.0, I13.2, I25.5, I42.0, I42.5-I42.9, I43, I50, P29.0 |
| **Asthma** | 493 | J45, J46 |

Diagnosis and external causes ICD codes in Western Australia’s Hospital Morbidity Data Collection. The ICD-9-CM codes commencing with “E” and ICD-10-AM codes commencing with “X” or “Y” are external cause codes. Diagnoses in the hospital data were coded using ICD-9-CM codes prior to 1 July 1999 and ICD-10-AM codes from 1 July 1999 onwards.

^1^ Codes used by O’Leary *et al*, with the exception of ICD codes for the offspring, such as fetal alcohol spectrum disorder, as these diagnoses are not recorded for stillborn children and could bias downwards any associations between perinatal death and alcohol and drug misuse [4].

^2^ Codes used by Derrington *et al* [5]

^3^ Syphilis, toxoplasmosis, rubella, cytomegalovirus and varicella zoster

^4^ Codes used by O’Donnell *et al*, excluding the alcohol and drug-related codes [6].

**2. Missing data**

Maternal height was missing for 9305/28119 (33%) of births. Additionally, for some women, the maternal heights listed on their children’s records were variable. To deal with these issues, we created a new height variable (“Imputation Step 1”), as follows:

- the height recorded for teenage mothers was retained, unless the mother ‘shrank’ with age;
- for births where the mother was at least 20 years, mothers were assigned their median ‘adult’ height;
- if a mother was recorded as losing height during her teenage years, or if her final teenage height exceeded her median adult height, her height for all teenage births was set to her median teenage height;
- if a mother’s median teenage height exceeded her median adult height, her height was set to her median height across all births; and
- where possible, the remaining missing heights were then set to the height for the birth closest in time. These included cases of multiple teenage births where at least one record was missing a height value or the mother had her height recorded for births as a teenager, but not for births as an adult.

For 6078 cases of missing maternal height, we were able to estimate height using the birth records of siblings, as outlined above. We then used multiple imputation to impute maternal height for the remaining 3227/28119 (11%) of births (“Imputation Step 2”) using the “mice” package in R 3.4.0 [7, 8].

The missing data was assumed to be missing at random (MAR). It is likely that missing maternal heights have a different distribution to observed heights, but this may be fully explained by year of birth. Prior to Imputation Step 1, only 10% of birth records for 1998 were missing maternal height, compared to 57% in 2008, suggesting changes in the collection of maternal height. Conditional on year of birth, it seems reasonable to assume there was no or little difference between observed and missing maternal heights. While the height of women with limited antenatal care may not have been measured, any difference in the heights of women with good antenatal care and limited care is likely to be small, after controlling for parity, maternal age, baby’s year of birth, maternal health, and health behaviours.

We specified a linear regression imputation model for maternal height, as a visual check of the data revealed an approximately normal distribution. Each of the three outcomes and each variable included in the three models presented in Supplementary Tables 3-5 were included as explanatory variables in the imputation model, with the exception that the powers for maternal age were initially different as powers 2 and 3 were selected for preterm birth using the complete case data and powers 1 and 3 were selected for perinatal death. We also initially included interactions that are not listed in the table, which were significant in the models for SGA based on complete case data, but not in those using imputed data (an alcohol and drug misuse interaction and a smoking and hypertension interaction).

We then analysed the imputed data and the best fractional polynomial for maternal age for the SGA model had powers 1 and 3 and for perinatal death -0.5 and 3. The interactions in the SGA model for between alcohol misuse and drug misuse and between smoking and hypertension were no longer significant. As a result, the imputation was redone with maternal age with power -0.5, 1 and 3 and the two non-significant interactions were dropped. We created 20 new complete datasets.

Supplementary Table 2 lists some maternal height statistics for the maternal height as recorded in the original data and after each of the imputation steps. The mean heights before and after imputation varied little, although the heights imputed in Step 2 were higher than in Step 1 and the original data, reflecting the increase in maternal height over time and the high proportion of missing values toward the end of 1998-2010. The imputed heights were less variable than the original data, as some extreme heights were replaced by median heights in Step 1.

**Supplementary Table 2: Comparison of original maternal heights and imputed maternal heights**

|  | Number of births | Mean | Standard deviation | Minimum | Maximum |
| --- | --- | --- | --- | --- | --- |
| **Original data** |  |  |  |  |  |
| Non-missing | 18814 | 163.1 | 6.4 | 132 | 193 |
| Missing | 9305 |  |  |  |  |
| **After Imputation Step 1** |  |  |  |  |  |
| Non-missing | 24892 |  |  |  |  |
| *Not imputed or changed in Step 1* | *11611* | *163.1* | *6.3* | *139* | *187* |
| *Changed or imputed in Step 1* | *13281* | *163.1* | *6.0* | *139* | *190* |
| Missing | 3227 |  |  |  |  |
| **After Imputation Step 2** |  |  |  |  |  |
| Non-missing | 28119 |  |  |  |  |
| *Not imputed in Step 2* | *24892* | *163.1* | *6.1* | *139* | *190* |
| *Imputed in Step 2* | *3227* | *163.6** | *6.1** | *138.0** | *191.8** |
| Missing | 0 |  |  |  |  |
| * Summary statistics are from all 20 imputed datasets. | | | | | |

Odds ratios for the imputed data and the complete case data can be seen in Supplementary Tables 6-8.

**3. Cross-classification to mothers and fathers**

Children can be cross-classified to mothers and fathers. To account for this, we used the method of Miglioretti and Heagerty [9]. We obtained regression coefficient covariance matrices from models with clustering by mothers only, clustering by fathers only, and clustering by each pairing of a mother and a father who were common parents of at least one child. For those children where the identity of the father was not known (eg the infant did not have a birth registration record) [10], we assumed each child had a different father. We then obtained a final matrix by subtracting the matrix from clustering by mother-father combinations from the sum of the matrices from clustering by mother only and by father only. However, these matrices were very similar to those obtained by clustering on the mother. We therefore present the results from clustering by the mother only.

**4. Odds ratios for all covariates in fully-adjusted models for poor birth outcomes**

A subset of the aORs for maternal smoking during pregnancy, alcohol misuse, drug misuse, and assault from the final models for SGA, preterm birth and perinatal death were reported in *Poor birth outcomes among Aboriginal Western Australians and smoking, alcohol, and substance misuse*. Supplementary Tables 3-5 show the adjusted odds ratios (aORs) for all covariates in the models.

**Supplementary Table 3: Adjusted odds ratios of being small for gestational age for 28,052 Aboriginal singleton infants born in Western Australia, 1998-2010**

| Factor | Odds ratio (95% CI) | P-value |
| --- | --- | --- |
| *Risk factors of interest* |  |  |
| Maternal smoking and drug abuse |  | <0.001* |
| Neither | 1.00 |  |
| Smoking only | 2.28 (2.12, 2.46) |  |
| Drug abuse only | 2.52 (2.00, 3.19) |  |
| Both | 2.82 (2.44, 3.25) |  |
| Alcohol abuse | 2.18 (1.84, 2.58) | <0.001 |
| Assault against mother | 1.61 (1.42, 1.81) | <0.001 |
| *Demographic factors* |  |  |
| Male infant | 1.10 (1.03, 1.17) | 0.006 |
| Parity |  | <0.001 |
| 0 | 1.00 |  |
| 1 | 0.68 (0.63, 0.74) |  |
| 2 or more | 0.62 (0.57, 0.67) |  |
| *Maternal health* |  |  |
| Maternal height (per cm increase) | 0.96 (0.95, 0.97) | <0.001 |
| Diabetes | 0.48 (0.40, 0.58) | <0.001 |
| Hypertension | 1.35 (1.21, 1.51) | <0.001 |
| Obesity | 0.47 (0.33, 0.67) | <0.001 |
| Gonorrhoea | 1.77 (1.26, 2.47) | <0.001 |
| Herpes | 0.60 (0.41, 0.89) | 0.01 |
| Other infections | 1.66 (1.12, 2.46) | 0.01 |
| Odds ratios were obtained from a model containing all of the above variables, including an interaction term for maternal smoking and drug abuse. "Other infections" refers to syphilis, toxoplasmosis, rubella, cytomegalovirus and/or varicella zoster. 67 cases of unknown gestational age were excluded from the sample of 28,119.  * P-values for maternal smoking, drug abuse and the interaction term were all <0.001. | | |

**Supplementary Table 4: Adjusted odds ratios of preterm birth for 28,119 Aboriginal singleton infants born in Western Australia, 1998-2010**

| Factor | Odds ratio (95% CI) | P-value |
| --- | --- | --- |
| *Risk factors of interest* |  |  |
| Maternal smoking | 1.26 (1.17, 1.36) | <0.001 |
| Drug misuse and vaginitis |  | <0.001* |
| Neither | 1 |  |
| Drug misuse only | 2.16 (1.88, 2.49) |  |
| Vaginitis only | 2.36 (2.06, 2.70) |  |
| Both | 2.88 (2.17, 3.81) |  |
| Alcohol misuse | 1.16 (0.95, 1.42) | 0.14 |
| Assault against mother | 1.40 (1.22, 1.60) | <0.001 |
| *Demographic factors* |  |  |
| Maternal age (reference age 25 years) |  | Power 1: <0.001 Power 3: <0.001 |
| 15 years | 1.40 (1.21, 1.62) |  |
| 20 years | 1.15 (1.08, 1.22) |  |
| 25 years | 1.00 |  |
| 30 years | 0.95 (0.91, 0.99) |  |
| 35 years | 1.02 (0.94, 1.12) |  |
| 40 years | 1.28 (1.06, 1.55) |  |
| Parity |  | <0.001 |
| 0 | 1.00 |  |
| 1 | 1.21 (1.09, 1.34) |  |
| 2 or more | 1.35 (1.20, 1.52) |  |
| Infant's year of birth (per year increase) | 1.01 (1.00, 1.02) | 0.02 |
| *Maternal health* |  |  |
| Maternal height (per cm increase) | 0.98 (0.97, 0.98) | <0.001 |
| Diabetes | 1.53 (1.33, 1.75) | <0.001 |
| Hypertension | 1.88 (1.69, 2.10) | <0.001 |
| Heart disease | 1.45 (1.04, 2.02) | 0.03 |
| Urinary tract infection | 1.11 (1.01, 1.23) | 0.04 |
| Group B streptococcus | 1.24 (1.07, 1.44) | 0.005 |
| Obesity | 1.28 (1.03, 1.59) | 0.02 |
| Mental health condition | 1.16 (1.01, 1.33) | 0.03 |
| Gonorrhoea | 1.68 (1.18, 2.40) | 0.004 |
| Odds ratios were obtained from a model containing all of the above variables, including an interaction term for drug misuse and vaginitis. Vaginitis also includes candida and trichomoniasis. Hypertension refers to pre-existing hypertension complicating pregnancy, pre-eclampsia, and eclampsia. Maternal age was modelled with a fractional polynomial with powers 1 and 3.  * P-values for drug misuse, vaginitis and the interaction term were all <0.001. | | |

**Supplementary Table 5: Adjusted odds ratios of perinatal death for 28,119 Aboriginal singleton infants born in Western Australia, 1998-2010**

| Factor | Odds ratio (95% CI) | P-value |
| --- | --- | --- |
| *Risk factors of interest* |  |  |
| Maternal smoking | 1.49 (1.23, 1.80) | <0.001 |
| Drug abuse | 1.06 (0.74, 1.54) | 0.75 |
| Alcohol abuse | 1.83 (1.16, 2.88) | 0.01 |
| Assault against mother | 0.90 (0.61, 1.34) | 0.61 |
| *Demographic factors* |  |  |
| Maternal age (reference age 25 years) |  | Power -0.5: 0.005 Power 3: 0.003 |
| 15 years | 1.60 (1.13, 2.27) |  |
| 20 years | 1.09 (1.00, 1.19) |  |
| 25 years | 1.00 |  |
| 30 years | 1.05 (0.98, 1.13) |  |
| 35 years | 1.26 (1.04, 1.55) |  |
| 40 years | 1.75 (1.15, 2.66) |  |
| *Maternal health* |  |  |
| Maternal height (per cm increase) | 0.98 (0.97, 1.00) | 0.05 |
| Diabetes | 1.49 (1.09, 2.05) | 0.02 |
| Urinary tract infection | 1.34 (1.06, 1.69) | 0.02 |
| Odds ratios were obtained from a model containing all of the above variables. Maternal age was modelled with a fractional polynomial with powers , 0.5 and 3. | | |

**5. Sensitivity analyses**

Supplementary Tables 6-8 contain the odds ratios obtained from identical models to those presented in Figure 2 of *Poor birth outcomes among Aboriginal Western Australians and smoking, alcohol, and substance misuse* and Supplementary Tables 3-5 with the following differences:

- complete case only. For these analyses, the sample was restricted to those with maternal height recorded on their own birth record;
- remoteness was included as an explanatory variable; and
- socioeconomic disadvantage was included as an explanatory variable.

**Supplementary Table 6: Comparison of adjusted odds ratios for small for gestational age using imputed and complete data, and additionally adjusting for remoteness or socioeconomic disadvantage**

| Factor | Odds ratio (95% CI) | P-value |
| --- | --- | --- |
| **Final model with missing maternal height imputed (n=28,052)** | |  |
| Maternal smoking and drug misuse |  | <0.001* |
| Neither | 1.00 |  |
| Smoking only | 2.28 (2.12-2.46) |  |
| Drug misuse only | 2.52 (2.00-3.19) |  |
| Both | 2.82 (2.44-3.25) |  |
| Alcohol misuse | 2.18 (1.84-2.58) | <0.001 |
| Assault against mother | 1.61 (1.42-1.81) | <0.001 |
| **Complete case analysis (n=18,785)** |  |  |
| Maternal smoking and drug misuse |  | <0.001* |
| Neither | 1.00 |  |
| Smoking only | 2.19 (2.00-2.40) |  |
| Drug misuse only | 2.55 (1.95-3.32) |  |
| Both | 2.70 (2.28-3.19) |  |
| Alcohol misuse | 2.13 (1.75-2.60) | <0.001 |
| Assault against mother | 1.54 (1.33-1.78) | <0.001 |
| **Adjusted for remoteness (n=28,052)** |  |  |
| Maternal smoking and drug misuse |  | <0.001* |
| Neither | 1.00 |  |
| Smoking only | 2.22 (2.06-2.39) |  |
| Drug misuse only | 2.80 (2.21-3.53) |  |
| Both | 3.09 (2.68-3.58) |  |
| Alcohol misuse | 2.06 (1.74-2.44) | <0.001 |
| Assault against mother | 1.50 (1.33-1.69) | <0.001 |
| **Adjusted for socioeconomic disadvantage (n=28,052)** |  |  |
| Maternal smoking and drug misuse |  | <0.001* |
| Neither | 1.00 |  |
| Smoking only | 2.22 (2.06-2.39) |  |
| Drug misuse only | 2.73 (2.16-3.44) |  |
| Both | 3.00 (2.60-3.47) |  |
| Alcohol misuse | 2.08 (1.76-2.47) | <0.001 |
| Assault against mother | 1.51 (1.34-1.71) | <0.001 |
| All models adjusted for maternal smoking, drug misuse, alcohol misuse, assault against the mother, infant sex, parity, maternal height, diabetes, hypertension (pre-existing hypertension complicating pregnancy, pre-eclampsia, and eclampsia), obesity, gonorrhoea, herpes, other infections (syphilis, toxoplasmosis, rubella, cytomegalovirus, and varicella zoster), and an interaction term between maternal smoking and drug misuse.  * P-values for maternal smoking, drug misuse, and the interaction term were all <0.001. | | |

**Supplementary Table 7: Comparison of adjusted odds ratios for preterm birth using imputed and complete data, and additionally adjusting for remoteness or socioeconomic disadvantage**

| Factor | Odds ratio (95% CI) | P-value |
| --- | --- | --- |
| **Final model with missing maternal height imputed (n=28,119)** | |  |
| Maternal smoking | 1.26 (1.17-1.36) | <0.001 |
| Drug misuse and vaginitis |  | <0.001* |
| Neither | 1.00 |  |
| Drug misuse only | 2.16 (1.88-2.49) |  |
| Vaginitis only | 2.36 (2.06-2.70) |  |
| Both | 2.88 (2.17-3.81) |  |
| Alcohol misuse | 1.16 (0.95-1.42) | 0.14 |
| Assault against mother | 1.40 (1.22-1.60) | <0.001 |
| **Complete case analysis (n=18,814)** |  |  |
| Maternal smoking | 1.29 (1.18-1.42) | <0.001 |
| Drug misuse and vaginitis |  | Drug and vaginitis: <0.001  Interaction: 0.007 |
| Neither | 1.00 |  |
| Drug misuse only | 2.15 (1.84-2.51) |  |
| Vaginitis only | 2.36 (2.03-2.75) |  |
| Both | 3.12 (2.26-4.31) |  |
| Alcohol misuse | 1.06 (0.84-1.33) | 0.65 |
| Assault against mother | 1.27 (1.08-1.49) | 0.003 |
| **Adjusted for remoteness (n=28,119)** |  |  |
| Maternal smoking | 1.25 (1.16-1.35) | <0.001 |
| Drug misuse and vaginitis |  | <0.001* |
| Neither | 1.00 |  |
| Drug misuse only | 2.22 (1.92-2.56) |  |
| Vaginitis only | 2.35 (2.05-2.70) |  |
| Both | 2.98 (2.24-3.95) |  |
| Alcohol misuse | 1.13 (0.92-1.38) | 0.23 |
| Assault against mother | 1.36 (1.18-1.57) | <0.001 |
| **Adjusted for socioeconomic disadvantage (n=28,119)** |  |  |
| Maternal smoking | 1.24 (1.15-1.34) | <0.001 |
| Drug misuse and vaginitis |  | <0.001* |
| Neither | 1.00 |  |
| Drug misuse only | 2.25 (1.96-2.60) |  |
| Vaginitis only | 2.37 (2.06-2.71) | <0.001 |
| Both | 3.03 (2.29-4.02) |  |
| Alcohol misuse | 1.13 (0.92-1.38) | 0.24 |
| Assault against mother | 1.35 (1.17-1.55) | <0.001 |
| All models adjusted for maternal smoking, drug misuse, alcohol misuse, assault against the mother, maternal age, parity, infant's year of birth, maternal height, diabetes, hypertension (pre-existing hypertension complicating pregnancy, pre-eclampsia, and eclampsia), heart disease, urinary tract infection, Group B streptococcus, obesity, mental health conditions, gonorrhoea, and an interaction between drug misuse and vaginitis.  * P-values for drug misuse, vaginitis, and the interaction term were all <0.001. | | |

**Supplementary Table 8: Comparison of adjusted odds ratios for perinatal death using imputed and complete data, and additionally adjusting for remoteness or socioeconomic disadvantage**

| Factor | Odds ratio (95% CI) | P-value |
| --- | --- | --- |
| **Final model with missing maternal height imputed (n=28,119)** | |  |
| Maternal smoking | 1.49 (1.23-1.80) | <0.001 |
| Drug misuse | 1.06 (0.74-1.54) | 0.75 |
| Alcohol misuse | 1.83 (1.16-2.88) | 0.01 |
| Assault against mother | 0.90 (0.61-1.34) | 0.61 |
| **Complete case analysis, all parities (n=18,814)** |  |  |
| Maternal smoking | 1.66 (1.32-2.09) | <0.001 |
| Drug misuse | 1.08 (0.71-1.64) | 0.71 |
| Alcohol misuse | 1.29 (0.72-2.31) | 0.40 |
| Assault against mother | 0.84 (0.53-1.34) | 0.47 |
| **Adjusted for remoteness (n=28,119)** |  |  |
| Maternal smoking | 1.45 (1.20-1.76) | <0.001 |
| Drug misuse | 1.17 (0.80-1.70) | 0.42 |
| Alcohol misuse | 1.72 (1.09-2.71) | 0.02 |
| Assault against mother | 0.85 (0.57-1.26) | 0.41 |
| **Adjusted for socioeconomic disadvantage (n=28,119)** |  |  |
| Maternal smoking | 1.45 (1.19-1.76) | <0.001 |
| Drug misuse | 1.15 (0.80-1.67) | 0.45 |
| Alcohol misuse | 1.72 (1.09-2.73) | 0.02 |
| Assault against mother | 0.85 (0.58-1.25) | 0.41 |
| All models adjusted for maternal smoking, drug misuse, alcohol misuse, assault against the mother, maternal age, maternal height, diabetes, and urinary tract infection. | | |

The population attributable fractions (PAFs) for each risk factor and each birth outcome are listed in Supplementary Table 9, along with the results from models which also included remoteness or socioeconomic disadvantage.

**Supplementary Table 9: Comparison of population attributable fractions (PAFs) and 95% confidence intervals (95% CI) for birth outcomes using final models and additionally adjusting for remoteness or socioeconomic disadvantage**

| **Factor** |  | **PAF (95% CI)** |  |
| --- | --- | --- | --- |
|  | **Final models** | **+ remoteness** | **+ socioeconomic disadvantage** |
| **Small for gestational age** |  |  |  |
| Maternal smoking | 28.1 (25.4, 30.9) | 27.3 (24.5, 30.1) | 27.2 (24.5, 30.1) |
| Drug misuse | 2.5 (1.6, 3.3) | 3.2 (2.4, 4.1) | 3.1 (2.2, 3.9) |
| Alcohol misuse | 2.7 (2.1, 3.3) | 2.5 (1.9, 3.2) | 2.5 (1.9, 3.2) |
| Assault against mother | 3.5 (2.5, 4.3) | 3.0 (2.1, 3.9) | 3.1 (2.1, 4.0) |
| All of above | 37.5 (34.9, 40.0) | 36.7 (34.2, 39.3) | 36.7 (34.1, 39.3) |
| **Preterm birth** |  |  |  |
| Maternal smoking | 9.4 (6.4, 12.3) | 9.1 (6.0, 12.0) | 8.9 (5.9, 11.8) |
| Drug misuse | 5.0 (3.9, 6.2) | 5.2 (4.0, 6.3) | 5.3 (4.1, 6.4) |
| Alcohol misuse | 0.5 (-0.2, 1.2) | 0.4 (-0.2, 1.1) | 0.4 (-0.3, 1.1) |
| Assault against mother | 2.5 (1.5, 3.5) | 2.3 (1.3, 3.3) | 2.3 (1.3, 3.3) |
| All of above | 16.5 (13.5, 19.0) | 16.1 (13.5, 19.1) | 16.0 (13.0, 18.5) |
| **Perinatal death** |  |  |  |
| Maternal smoking | 18.6 (9.6, 26.6) | 17.6 (8.3, 26.1) | 17.6 (8.4, 26.0) |
| Drug misuse | 0.5 (-2.1, 3.4) | 1.1 (-1.4, 4.2) | 1.1 (-1.4, 4.1) |
| Alcohol misuse | 2.6 (0.5, 4.7) | 2.4 (0.3, 4.5) | 2.4 (0.3, 4.5) |
| Assault against mother | -0.8 (-4.0, 2.2) | -1.4 (-4.4, 1.7) | -1.4 (-4.4, 1.7) |
| All of above | 20.2 (11.4, 28.1) | 19.3 (10.0, 27.2) | 19.2 (9.9, 27.1) |
| The final model for small for gestational age adjusted for maternal smoking, drug misuse, alcohol misuse, assault against the mother, infant sex, parity, maternal height, diabetes, hypertension, obesity, gonorrhoea, herpes, and other infections. The final model for preterm birth adjusted for maternal smoking, drug misuse, alcohol misuse, assault against the mother, maternal age, parity, infant's year of birth, maternal height, diabetes, hypertension, heart disease, urinary tract infection, Group B streptococcus, obesity, mental health conditions, and gonorrhoea. The final model for perinatal death adjusted for maternal smoking, drug misuse, alcohol misuse, assault against the mother, maternal age, maternal height, diabetes, and urinary tract infection. Results in the third column are from models that also included remoteness and results in the fourth column are from models that also included socioeconomic disadvantage. | | | |

**6. Subgroup analyses**

Supplementary Tables 10-12 contain the odds ratios obtained from identical models to those in Supplementary Tables 3-5, stratified by parity.

**Supplementary Table 10: Adjusted odds ratios for small for gestational age, stratified by parity**

| Factor | Odds ratio (95% CI) | P-value |
| --- | --- | --- |
| **Nulliparous births (n=8,613)** |  |  |
| Maternal smoking and drug misuse |  | <0.001* |
| Neither | 1.00 |  |
| Smoking only | 2.22 (1.98-2.50) |  |
| Drug misuse only | 2.97 (2.02-4.39) |  |
| Both | 3.01 (2.38-3.82) |  |
| Alcohol misuse | 2.19 (1.53-3.15) | <0.001 |
| Assault against mother | 1.91 (1.52-2.41) | <0.001 |
| **Parity 1 (n=6,884)** |  |  |
| Maternal smoking and drug misuse |  | Smoking and drug: <0.001 Interaction: 0.04 |
| Neither | 1.00 |  |
| Smoking only | 2.28 (1.96-2.64) |  |
| Drug misuse only | 2.56 (1.56-4.20) |  |
| Both | 3.17 (2.36-4.27) |  |
| Alcohol misuse | 2.09 (1.44-3.03) | <0.001 |
| Assault against mother | 1.43 (1.11-1.84) | 0.005 |
| **Parity 2 or more (n=12,555)** |  |  |
| Maternal smoking and drug misuse |  | <0.001* |
| Neither | 1.00 |  |
| Smoking only | 2.33 (2.06-2.62) |  |
| Drug misuse only | 2.23 (1.58-3.16) |  |
| Both | 2.57 (2.07-3.19) |  |
| Alcohol misuse | 2.25 (1.80-2.80) | <0.001 |
| Assault against mother | 1.56 (1.32-1.83) | <0.001 |
| All models adjusted for maternal smoking, drug misuse, alcohol misuse, assault against the mother, infant sex, maternal height, diabetes, hypertension (pre-existing hypertension complicating pregnancy, pre-eclampsia, and eclampsia), obesity, gonorrhoea, herpes, other infections (syphilis, toxoplasmosis, rubella, cytomegalovirus, and varicella zoster), and an interaction term between maternal smoking and drug misuse.  * P-values for maternal smoking, drug misuse, and the interaction term were all <0.001. | | |

**Supplementary Table 11: Adjusted odds ratios for preterm births, stratified by parity**

| Factor | Odds ratio (95% CI) | P-value |
| --- | --- | --- |
| **Nulliparous births (n=8,636)** |  |  |
| Maternal smoking | 1.12 (0.98-1.29) | 0.10 |
| Drug misuse and vaginitis |  | Drug and vaginitis: <0.001  Interaction: 0.05 |
| Neither | 1.00 |  |
| Drug misuse only | 2.18 (1.69-2.79) |  |
| Vaginitis only | 1.61 (1.26-2.05) |  |
| Both | 1.81 (1.01-3.25) |  |
| Alcohol misuse | 1.54 (1.02-2.33) | 0.04 |
| Assault against mother | 1.31 (0.98-1.74) | 0.07 |
| **Parity 1 (n=6,898)** |  |  |
| Maternal smoking | 1.35 (1.16-1.57) | <0.001 |
| Drug misuse and vaginitis |  | Drug and vaginitis: <0.001  Interaction: 0.53 |
| Neither | 1.00 |  |
| Drug misuse only | 2.00 (1.50-2.68) |  |
| Vaginitis only | 2.35 (1.78-3.08) |  |
| Both | 3.79 (2.16-6.65) |  |
| Alcohol misuse | 1.16 (0.76-1.76) | 0.49 |
| Assault against mother | 1.58 (1.22-2.06) | <0.001 |
| **Parity 2 or more (n=12,585)** |  |  |
| Maternal smoking | 1.31 (1.17-1.46) | <0.001 |
| Drug use and vaginitis |  | Drug and vaginitis: <0.001 Interaction: 0.001 |
| Neither | 1.00 |  |
| Drug misuse only | 2.24 (1.83-2.73) |  |
| Vaginitis only | 3.09 (2.53-3.77) |  |
| Both | 3.28 (2.18-4.93) |  |
| Alcohol | 1.06 (0.81-1.39) | 0.65 |
| Assault against mother | 1.35 (1.12-1.62) | 0.001 |
| All models adjusted for maternal smoking, drug misuse, alcohol misuse, assault against the mother, maternal age, infant's year of birth, maternal height, diabetes, hypertension (pre-existing hypertension complicating pregnancy, pre-eclampsia, and eclampsia), heart disease, urinary tract infection, Group B streptococcus, obesity, mental health conditions, gonorrhoea, and an interaction between drug misuse and vaginitis.  * P-values for drug misuse, vaginitis, and the interaction term were all <0.001. | | |

**Supplementary Table 12: Adjusted odds ratios for perinatal death, stratified by parity**

| Factor | Odds ratio (95% CI) | P-value |
| --- | --- | --- |
| **Nulliparous births (n=8,636)** |  |  |
| Maternal smoking | 1.23 (0.87-1.74) | 0.24 |
| Drug misuse | 0.85 (0.42-1.75) | 0.67 |
| Alcohol misuse | 3.57 (1.53-8.34) | 0.003 |
| Assault against mother | 0.60 (0.24-1.53) | 0.29 |
| **Parity 1 (n=6,898)** |  |  |
| Maternal smoking | 1.58 (1.05-2.37) | 0.03 |
| Drug misuse | 0.58 (0.21-1.63) | 0.30 |
| Alcohol misuse | 2.11 (0.78-5.69) | 0.14 |
| Assault against mother | 0.91 (0.40-2.06) | 0.82 |
| **Parity 2 or more (n=12,585)** |  |  |
| Maternal smoking | 1.61 (1.22-2.14) | <0.001 |
| Drug misuse | 1.36 (0.87-2.14) | 0.18 |
| Alcohol misuse | 1.38 (0.73-2.59) | 0.32 |
| Assault against mother | 0.99 (0.61-1.61) | 0.97 |
| All models adjusted for maternal smoking, drug misuse, alcohol misuse, assault against the mother, maternal age, maternal height, diabetes, and urinary tract infection. | | |

**References**

1.	Australian Institute of Health and Welfare and University of New South Wales. 2017. Data Collection Overview, Western Australian Midwives' Notification System (WA PDC). http://maternitymatrix.aihw.gov.au/Pages/CollDetails.aspx?DataCollID=14. Accessed 31 January 2017.

2.	Blair E, Liu Y, Cosgrove P. Choosing the best estimate of gestational age from routinely collected population-based perinatal data. Paediatr Perinat Epidemiol. 2004;18:270-276.

3.	Dobbins TA, Sullivan EA, Roberts CL, Simpson JM. Australian national birthweight percentiles by sex and gestational age, 1998-2007. Med J Aust. 2012;197:291-294.

4.	O'Leary CM, Watson L, D'Antoine H, Stanley F, Bower C. Heavy maternal alcohol consumption and cerebral palsy in the offspring. Dev Med Child Neurol. 2012;54:224-230.

5.	Derrington TM, Bernstein J, Belanoff C, Cabral HJ, Babakhanlou-Chase H, Diop H, et al. Refining Measurement of Substance Use Disorders Among Women of Child-Bearing Age Using Hospital Records: The Development of the Explicit-Mention Substance Abuse Need for Treatment in Women (EMSANT-W) Algorithm. Matern Child Health J. 2015.

6.	O'Donnell M, Anderson D, Morgan VA, Nassar N, Leonard HM, Stanley FJ. Trends in pre-existing mental health disorders among parents of infants born in Western Australia from 1990 to 2005. Med J Aust. 2013;198:485-488.

7.	van Buuren S, Groothuis-Oudshoorn K. mice: Multivariate imputation by chained equations in R. J Stat Softw. 2011;45.

8.	van Buuren S. Flexible imputation of missing data. Boca Raton (FL): CRC press; 2012.

9.	Miglioretti DL, Heagerty PJ. Marginal modeling of nonnested multilevel data using standard software. Am J Epidemiol. 2007; 165:453-463.

10.	Gibberd AJ, Simpson JM, Eades SJ. No official identity: a data linkage study of birth registration of Aboriginal children in Western Australia. Aust N Z J Public Health. 2016; 40:388-394.
